# Multiplexed nanopore amplicon sequencing to distinguish recrudescence from new infection in antimalarial drug trials

**DOI:** 10.1101/2024.09.11.612449

**Authors:** Aurel Holzschuh, Anita Lerch, Christian Nsanzabana

**Affiliations:** Swiss Tropical and Public Health Institute, Allschwil, Switzerland; University of Basel, Basel, Switzerland; Department of Biological Sciences, Eck Institute for Global Health, University of Notre Dame, Indiana, USA

**Keywords:** Malaria, *Plasmodium falciparum*, genomics, amplicon sequencing, nanopore sequencing, clinical trial, therapeutic efficacy study, multiplex PCR, haplotype, genetic diversity

## Abstract

**Background:** The assessment of antimalarial drug efficacy against *Plasmodium falciparum* requires PCR correction to distinguish recrudescence from new infections by comparing parasite genotypes before treatment and in recurrent infections. Nanopore sequencing offers a low-cost, portable, scalable, and rapid alternative to traditional methods, supporting the expansion and decentralization of sequencing in endemic, resource-limited settings, potentially providing rapid PCR-corrected drug failure estimates.

**Methods:** We optimized a multiplexed AmpSeq panel targeting six microhaplotypes for high and uniform coverage. We assessed sensitivity and specificity for detecting minority clones in polyclonal infections and evaluated genetic diversity across the microhaplotype markers. We used mixtures of four *P. falciparum* laboratory strains at different ratios and 20 paired patient samples from a clinical trial. A custom bioinformatics workflow was used to infer haplotypes from polyclonal infections, including minority clones, with defined cut-off criteria for accurate haplotype calling.

**Findings:** The nanopore AmpSeq assay achieved uniform and high read coverage across all six microhaplotype markers (median coverage: 12,989× to 15,440× for laboratory strain mixtures and 7,011× to 11,600× for patients’ samples, respectively). We found high sensitivity in detecting minority clones (up to 50:1:1:1 in the 3D7:K1:HB3:FCB1 laboratory strain mixtures) and high specificity with less than 0.01% of all reads being false-positive haplotypes. Genetic diversity in the markers used was high (H_E_ ≥ 0.98 and up to 31 unique haplotypes in 20 paired samples with *cpmp*), and concordant results in classifying new infections and recrudescence across all markers used were observed in 18 (90%) of 20 paired samples.

**Interpretation:** Our study demonstrates the feasibility of nanopore AmpSeq for distinguishing recrudescence from new infections in clinical trials.

**Funding:** Swiss Tropical and Public Health Institute.

## INTRODUCTION

The emergence of drug-resistant *Plasmodium falciparum* parasites is a significant public health concern that threatens global efforts to control and eliminate malaria. For the past two decades, the standard treatment for uncomplicated *P. falciparum* malaria has been artemisinin-based combination therapy (ACT). Delayed clearance following treatment with artemisinin derivatives, also referred to as artemisinin partial resistance (ART-R), was first described in 2008 ^1^ and was initially confined to Southeast Asia. However, recent independent emergence of ART-R in several regions of East Africa ^2–5^ is challenging ACT efficacy and malaria control in sub-Saharan Africa. This underscores the urgent need to improve genomic surveillance of ART-R across the continent and strengthen antimalarial drug efficacy surveillance through routine therapeutic efficacy studies (TES). Widespread ART-R combined with resistance to partner drugs (i.e., ACT failure) in sub-Saharan Africa would have a catastrophic public health impact, as the region bears the heaviest brunt of malaria burden ^6^. Regular reporting of drug efficacy across sites in Africa is essential, particularly in areas with high reported *P. falciparum kelch13* (*pfk13*) mutations, the key mediator of ART-R ^7^.

Antimalarial drug efficacy is evaluated through clinical and parasitological assessments. However, the cure rates obtained at the end of the follow-up period require PCR-correction to distinguish recrudescence (treatment failure) from new infection, especially in high-transmission settings where patients frequently acquire new infections after treatment ^8^. Different methods and molecular markers have been employed by different laboratories, along with various algorithms for interpreting genotyping results ^9,10^, making it challenging to compare drug efficacy estimates over time and space. The World Health Organization (WHO) currently recommends genotyping three size polymorphic markers - *P. falciparum* merozoite surface protein 1 and 2 (*pfmsp1* and *pfmsp2*) and microsatellites - using capillary electrophoresis ^8^.

Recent advances in next-generation sequencing (NGS) techniques have facilitated the development of targeted amplicon sequencing (AmpSeq) protocols of highly polymorphic markers for genotyping malaria parasites in clinical trials ^11–13^. By targeting short, highly diverse loci with multiple single nucleotide polymorphisms (SNPs), such microhaplotypes can greatly improve discrimination between infections ^14–18^. Detection of minority clones is possible at frequencies as low as 1% in polyclonal infections ^11,12^, and data can be analyzed using inferential methods considering allele sharing probabilities based on population frequencies ^19,20^. Despite its advantages, AmpSeq has not yet been used as the primary genotyping method in clinical trials. Small multiplexed AmpSeq panels of 3 to 5 highly diverse microhaplotypes are potentially more informative than traditional length-polymorphic or allele-specific genotyping techniques, while also being more cost-effective ^11,12,16^.

Recently, WHO has emphasized strengthening surveillance capabilities in Africa by increasing laboratory and technical capacity, and expanding data collection on antimalarial drug resistance and efficacy ^21^. Oxford Nanopore Technologies (ONT) offers a low-cost, portable, and scalable alternative to traditional sequencing methods, potentially addressing the need to expand and decentralize sequencing capacity in resource-limited settings. Workflows with ONT are straightforward, offer fast turnaround times, and are more easily deployed in endemic settings than other platforms ^22–24^. Thus, nanopore AmpSeq could provide rapid PCR-corrected estimates of drug efficacy, for example through an unplanned TES after rapid emergence of drug resistant parasites.

While previous studies primarily focused on malaria genomic surveillance, few studies have used nanopore AmpSeq to successfully explore within-host diversity and detection of minority clones in polyclonal infections ^22^. Here, we demonstrate the suitability of nanopore AmpSeq for genotyping *P. falciparum* to distinguish recrudescence from new infection in clinical trials, using the current latest ONT chemistry (kit 14/R10.4.1 flow cells). We optimized a 6-plex of short microhaplotypes (∼200 bp) for high and uniform coverage. We leveraged laboratory strain mixtures and paired patient samples, previously employed for a comprehensive comparison of five different genotyping methods, including AmpSeq using Illumina, to distinguish recrudescence from new infection ^13^.

## METHODS

### Laboratory strain mixtures

Laboratory *P. falciparum* strain mixtures (3D7, K1, HB3 and FCB1) were prepared as previously described ^13^. Twenty-four different ratios ranging from 1:1:1:1 to 1:100:100:100 were used for this study, with the concentration of the minority clone always being 10 parasites/μL (Supplementary Table 1).

### Patient samples

The samples were collected from a clinical trial evaluating the efficacy and safety of cipargamin (KAE609) in a randomized, dose-escalation, Phase II study in adults with uncomplicated *P. falciparum* malaria in high-transmission settings in sub-Saharan Africa. Details of the study protocol and patient recruitment have been published elsewhere ^25^. Twenty paired samples with recurrent infections were used in this study. The samples had been previously analyzed to compare different genotyping techniques, including AmpSeq using Illumina ^13^.

### Multiplex PCR amplification of microhaplotypes

We used a 6-plex PCR panel targeting highly polymorphic microhaplotypes (*ama1, celtos, cpmp, cpp, csp*, and *surfin1*.*1*; Supplementary Table 2) ^22^, with optimized primer concentrations and thermocycling conditions to ensure uniform amplification across all targets. The final 25 μL multiplex PCR reaction consisted of 0.5 μL combined primer mix, 12.5 μL KAPA HiFi HotStart ReadyMix (2X), 8 μL nuclease-free water, and 4 μL DNA template. Final primer pool concentrations and PCR cycling conditions are listed in Supplementary Table 3 and 4.

### Nanopore library preparation and sequencing

Library preparation was performed using ONT kit SQK-NBD114.96 following the manufacturer’s protocol (version NBA_9170_v114_revL_15Sep2022) with some modifications. Briefly, in the “end-prep” step, the incubation time was increased from 20ºC for 5 min and 65ºC for 5 min to 15 min for each step. End-prepped DNA was bead-purified using 1.6X ratio AMPure XP beads (Beckman Coulter) and resuspended in 15 μL nuclease-free water. For the “native barcoding ligation”, 3.75 μL end-prepped DNA was used instead of 0.75 μL, the incubation time was increased from 20 min to 30 min, and barcoded samples were bead-purified using 1X ratio AMPure XP beads instead of 0.4X. For the “adapter ligation and clean-up” step, the incubation time was increased from 20 min to 30 min and the pooled barcoded samples were bead-purified using 50 μL AMPure XP beads instead of 20 μL. Short fragment buffer was used for the wash steps. The final pooled library was quantified on a Qubit fluorometer (ThermoFisher), diluted in elution buffer (ONT) to a total of ∼250 fmol, and loaded onto R10.4.1 flow cells. Sequencing was performed on a MinION Mk1C instrument (ONT) with MinKNOW software (distribution version 23.07.12, core version 5.7.5, and configuration version 5.7.11). Twenty-four control mixtures in triplicate (*n*=72) and twenty paired patient samples (*n*=40) were sequenced in two MinION runs. At least one negative control, consisting of nuclease-free water, was included in each run and underwent the entire workflow, including PCR and nanopore library preparation. A positive control (*FCB1*) was only included for the paired patient samples. An overview of the entire workflow is shown in Supplementary Figure 1.

### Bioinformatics

Raw nanopore data (*.pod5 files) were simplex basecalled with *dorado* (v0.7.0; https://github.com/nanoporetech/dorado) using the super-accurate (sup) model (dna_r10.4.1_e8.2_400bps_sup@v5.0.0) with the --no-trim flag set. To ensure high-quality data for haplotype inference, the minimum q-score for passing reads was set to a stringent value of 20 to minimize erroneous reads (--min-qscore 20 flag; accuracy of ≥99%), as previously described ^22^. Raw reads passing quality filtering criteria were then demultiplexed by barcode using dorado (v0.7.0) for native barcodes with the --barcode-both-ends flag set, i.e., double-ended demultiplexing to reduce false positives. A read summary was created using *dorado* (v0.7.0) with the summary command from all basecalled data, i.e., without setting the --min-qscore 20 flag, and quality metrics were evaluated with NanoStat using the --summary option. Basecalling and demultiplexing were performed on a high-performance computing (HPC) cluster at the sciCORE scientific computing center at University of Basel (http://scicore.unibas.ch/).

From the resulting fastq files, haplotypes were inferred using the R packages *HaplotypR* (v0.5.0; https://github.com/lerch-a/HaplotypR) ^11^ and *DADA2* (v1.26.0) ^26^ as previously described ^22^. Briefly, each sample was demultiplexed by marker using demultiplexByMarkerMinION() from *HaplotypR*. Reads were then filtered to remove reads with ambiguous base calls (e.g., N) and incorrect sequence length, and haplotypes were infered using the learnError() and dada() functions from *DADA2*. Read counts per haplotype were extracted with makeSequenceTable() and all haplotype calls were filtered with removeBimeraDenovo() to remove chimeric haplotypes. To remove false-positive haplotypes the following cut-off criteria were implemented: (1) each marker required a minimum of 100 total reads per sample, (2) haplotype calls required a minimum coverage of ≥50 reads per haplotype, and (3) a within-sample allele frequency (WSAF) of ≥1%. Cut-offs were defined based on the control mixtures to maximize sensitivity and reduce any false-positive haplotype calls. An overview of the bioinformatics workflow is shown in Supplementary Figure 2.

### Data analysis

The limit of detection (LOD) of minority clones was defined as the ratio of laboratory strains having at least two out of three replicates positive. Two algorithms were used to classify genotyping results for the six microhaplotype markers. The WHO/MMV algorithm defines sample pairs as recrudescent when all markers used show recrudescence ^8^, while the 2/3 algorithm classifies sample pairs as recrudescent when the majority of markers used (i.e., 4/6) show recrudescence ^27^. Allelic frequency was assessed by calculating the number of times, each haplotype occurred in all samples divided by the number of times all haplotypes occurred in all samples. Genetic diversity of the microhaplotypes in the twenty paired patient samples was assessed by expected heterozygosity (H_E_) using the equation 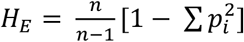 where *n* = the number of samples analyzed, *p*_*i*_ = the allele frequency of the *i*^th^ allele in the population. Multiplicity of infection (MOI) was defined as the highest number of alleles detected by any of the 6 microhaplotype loci. Pairwise genetic relatedness analysis was performed using *dcifer* (v1.2.0) with default settings ^20^. The concordance between markers was assessed in clinical samples, and agreement between markers sequenced by Illumina and ONT was visualized using the UpSetR package in R ^28^.

## RESULTS

For both MinION runs, reads from triplicates of control mixtures (*n*=72) and paired patient samples (*n*=40) had comparable q-scores, with >55% of reads being ≥Q20 (Figure 1a and c). Characteristics of both nanopore sequencing runs are shown in Supplementary Table 5. After several rounds of PCR optimizations and primer concentration (Supplementary Figure 3, Supplementary Table 6), we achieved high and uniform coverage across all amplicons in both, the control mixtures (Figure 1b) and paired patient samples (Figure 1d). After filtering reads with Q Score ≤Q20, the median coverage across triplicates of the 24 different control mixtures (*n*=72) was 84,993× per sample (interquartile range (IQR): 67,126× – 103,713×). The median coverage across all amplicons exceeded 10,000× per sample (range: 12,989× to 15,440× median coverage; Figure 1b). No sample had amplicons with coverage below 1,000×. Median coverage across all 20 paired patient samples was 54,942× (IQR: 39,548× – 73,075×). As with the control mixtures, reads were evenly distributed, with high median read coverage across amplicons (range: 7,011× to 11,600× median coverage, Figure 1d). One paired patient sample (S09) did not successfully amplify marker *cpp*. The median fold-difference in coverage between the amplicons was 1.4 (range: 1.1–10.8; IQR: 1.3–1.6) for control mixtures and 1.8 (range: 1.2–65.8; IQR: 1.4–2.1) for patient samples. Negative controls generated between 152 and 1,415 reads, but no haplotypes were called.

**Figure 1.**
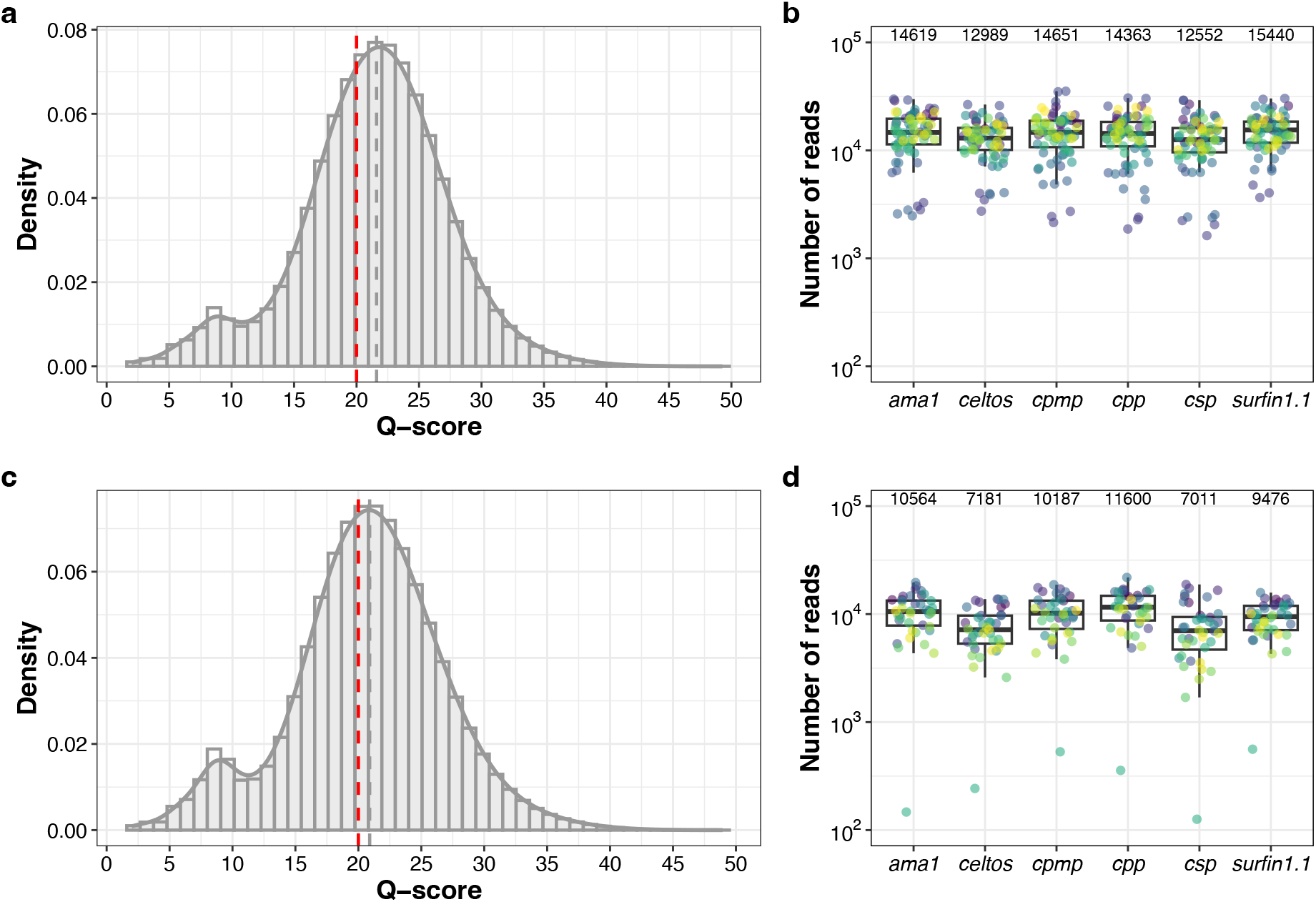
Quality metrics and read coverage profiles of two MinION sequencing runs. Distribution of q-scores (*dorado* v0.7.0) for the control mixtures (*n*=72; **a**) and paired patient samples (*n*=40; **c**). The dotted red line indicates the minimum q-score of Q20 for passing reads and the dotted grey line indicates the median q-scores. Median q-scores are 21.6 (99.3% accuracy) (**a**) and 20.9 (99.2% accuracy) (**c**), respectively. Coverage profiles of amplicon targets for the control mixtures (**b**) and paired patient samples (**d**), sequenced with kit 14 chemistry/R10.4.1 flow cells. Different colors represent different samples. The y-axis shows number of reads (log10) covering each amplicon target per sample for each of the two MinION runs. In both figures, positive and negative controls were excluded. Median coverage for all amplicons is indicated on top of the graph. The box bounds the IQR divided by the median, and Tukey-style whiskers extend to a maximum of 1.5 × IQR beyond the box.

### Specificity and limit of detection in control mixtures

The specificity of haplotype calls was high (Supplementary Figure 4). In control samples consisting of only one of the four *P. falciparum* strains (samples S01-S04), contaminating haplotypes (haplotypes from other samples of the same sequencing run) were found in only four instances, with a median of two reads per haplotype (range: 2–4). False-positive haplotypes (amplification or sequencing errors) were detected in 20 instances, with a median of 17 reads per haplotype (range: 6–214; IQR: 8–43). Overall, only 0.01% of total reads were contaminating or false positive haplotypes (833 of 6,170,352). All false-positive haplotype were below 1% WSAF. Thus, a conservative cut-off of ≥50 reads per haplotype and a WSAF of ≥1% were used to minimize false positives. When applying WSAF cut-offs below 1% (but keeping ≥50 reads per haplotype), false-positive haplotypes remained rare (Figure 2) and were only found in a single replicate (sample S11) for marker *surfin1*.*1*, present at frequencies of 0.53%, 0.72%, and 0.94%.

**Figure 2.**
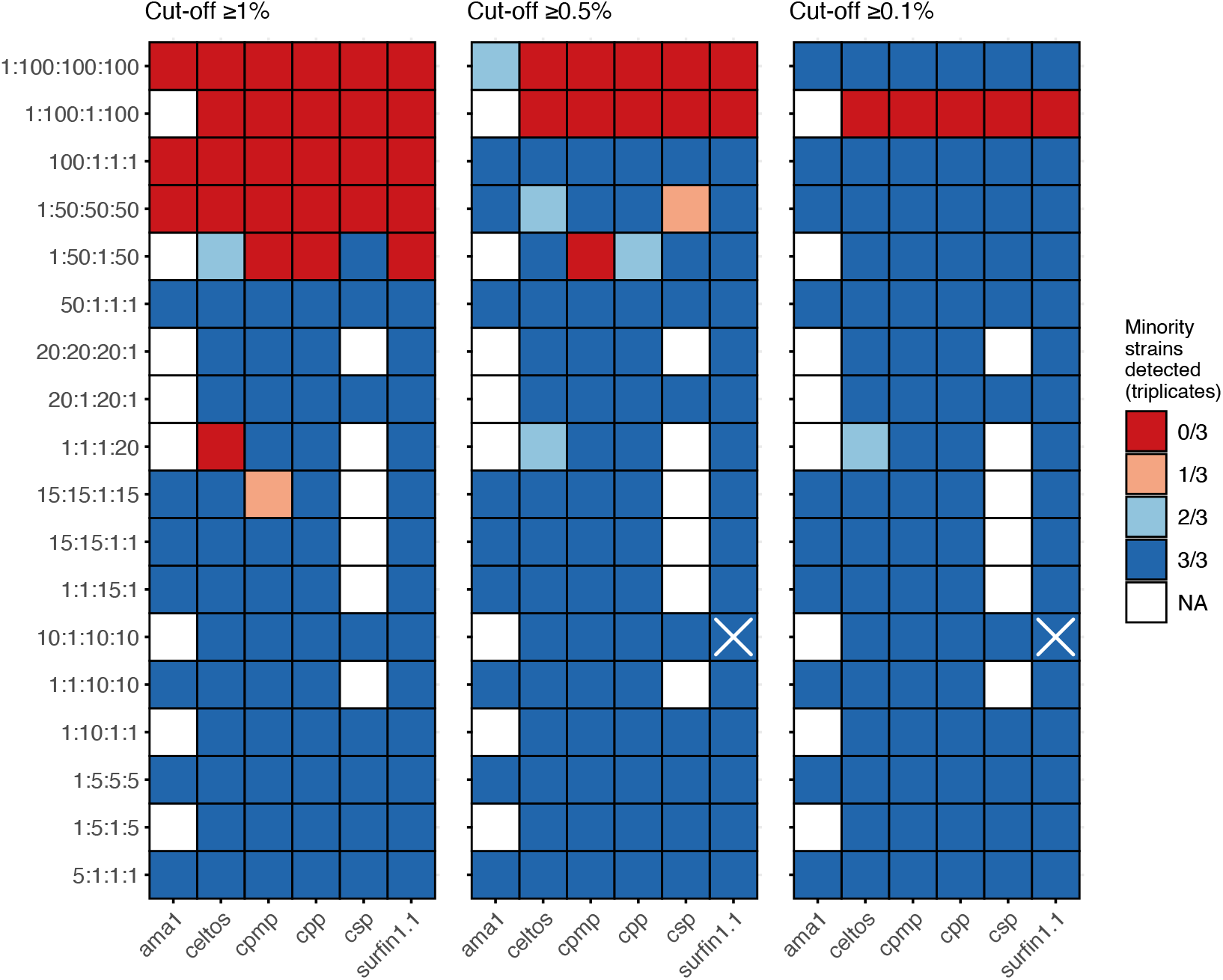
Limit of detection (LOD) for minority clones in control mixtures. WSAF cut-offs of ≥1% (left), ≥0.5% (middle), and ≥0.1% (right) are shown. Every control mixture was sequenced in triplicate and colors indicate in how many replicates the minority clones were detected at each WSAF cut-off. White cross indicates the detection of false-positive haplotypes above the cut-offs applied in one replicate (present at 0.94%, 0.72%, and 0.53%). The LOD for *ama1* and *csp* could not be assessed for all strain mixtures, as two strains had identical alleles (FCB1 and K1 for *ama1*; FCB1 and HB3 for *csp*; indicated in white color). Y-axis: *P. falciparum* laboratory strains mix ratios [3D7:K1:HB3:FCB1], with the concentration of the minority clone always being 10 parasites/μL. A minimum of 100 total reads per sample and a minimum coverage of ≥50 reads per haplotype were applied as described in the methods.

The LOD for minority clones was consistent across different markers (Figure 2). For two microhaplotypes, the LOD could not be determined for all four control mixtures because two strains had identical alleles for *ama1* (FCB1 and K1) and *csp* (FCB1 and HB3), respectively. Assessment for minority clone detection for these markers could only be done for two strains (3D7 and HB3 for *ama1*; 3D7 and K1 for *csp*) when one strain was the major clone and one the minor clone. At a WSAF of ≥1%, all six markers demonstrated the ability to detect the minority clone (10 parasites/μL) when the majority clones were at a 50-fold higher concentration (50:1:1:1), corresponding to WSAF of the minority clones of 1.9%. Intra-assay reproducibility was high, with the minority clone detected in all triplicates in all samples. Lowering the WSAF to ≥0.5% and ≥0.1% enabled detection of the minority clone in all triplicates, even when majority clones were at a 100-fold higher concentration (Figure 2).

### Haplotype diversity, and allelic frequency in paired patient samples

Diversity, defined as the number of distinct haplotypes in the patient samples, was high but varied among the six microhaplotypes (Figure 3, Table 1). Marker *cpmp* had 31 distinct haplotypes, whereas *surfin1*.*1* had the lowest number of distinct haplotypes (16). The number of SNPs for each marker ranged from 13 for *celtos* and *surfin1*.*1* to 38 for *cpmp* (Table 1). Allele frequency from patient samples varied across the microhaplotype markers (Figure 3). *Cpmp* did not have any haplotypes with an allelic frequency >10%, while *celtos* had only one haplotype at a relatively high frequency of 20% (Figure 3). *Ama1, cpp* and *csp* each had two haplotypes with an allelic frequency of >10%, without any clearly dominant haplotype, while *surfin1*.*1* had three dominant haplotypes, accounting for >50% of all *surfin1*.*1* sequence reads (Figure 3). The discriminatory power of the six markers, as estimated by expected heterozygosity (H_E_), was very high, with H_E_ ≥0.9 except for *surfin1*.*1* with an H_E_ < 0.9 (Table 1). Estimates of MOI were comparable across all markers, showing good concordance of mean MOI (*P* > 0.05 for all comparisons using Bonferroni-adjusted *t-test*, Table 1).

**Figure 3.**
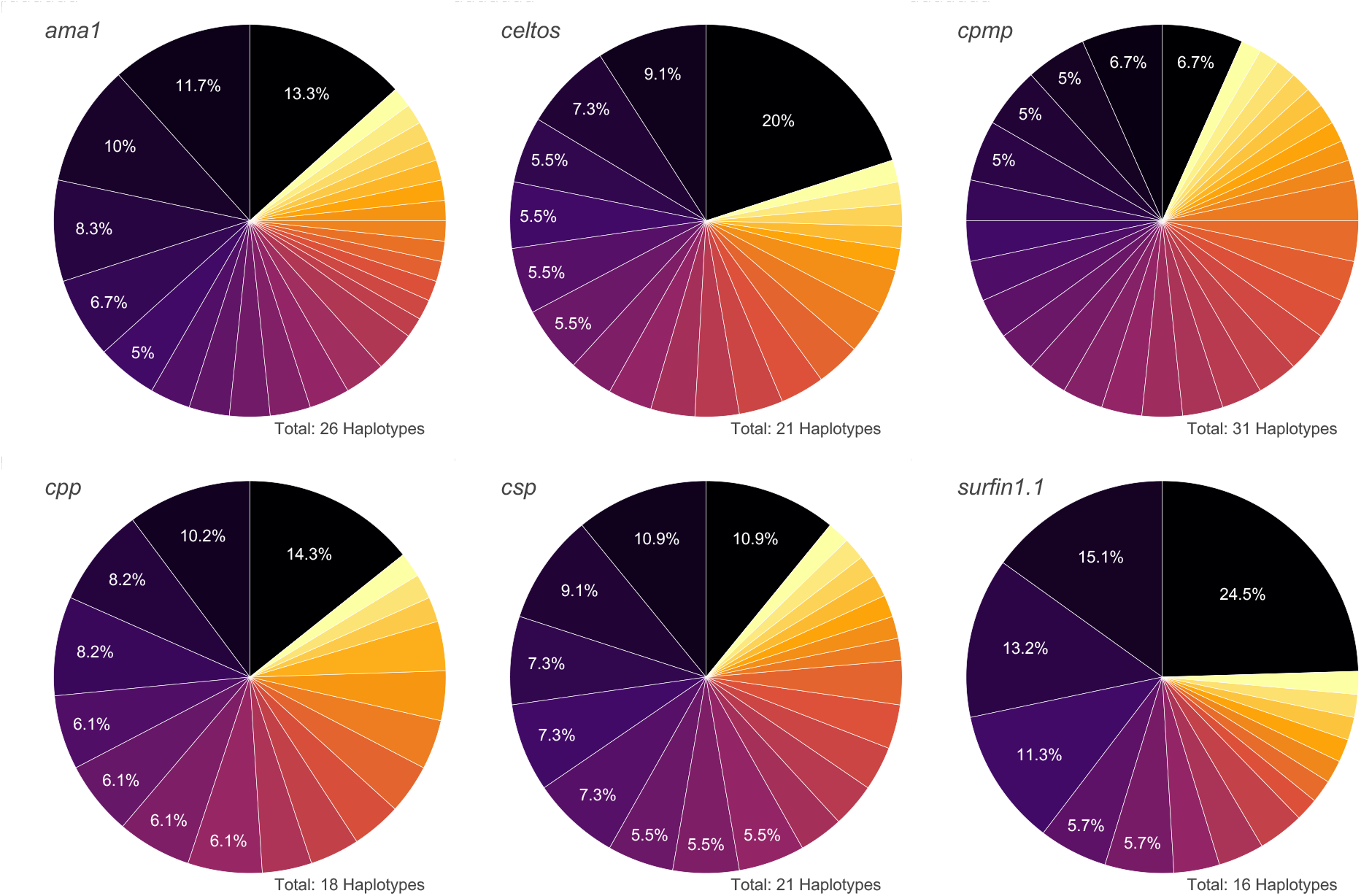
Diversity of the six microhaplotypes in all 20 paired patient samples. Each pie chart shows the distribution of genotypes for each marker across the patient sample pool (*n*=40). The total number of haplotypes found is shown at the bottom right of each microhaplotype pie chart. Haplotype frequencies greater than 5% are indicated.

**Table 1.** Diversity metrics of six microhaplotype marker in 20 paired patient samples using nanopore AmpSeq.

| Marker | Number of haplotypes | Number of SNPs | Expected heterozygosity ($H_E$) | Mean MOI (range) |
| --- | --- | --- | --- | --- |
| *ama1* | 26 | 22 | 0.949 | 1.50 (1 – 6) |
| *celtos* | 21 | 13 | 0.938 | 1.38 (1 – 6) |
| *cpmp* | 31 | 38 | 0.979 | 1.50 (1 – 6) |
| *cpp* | 18 | 24 | 0.947 | 1.28 (1 – 4) |
| *csp* | 21 | 14 | 0.952 | 1.38 (1 – 4) |
| *surfin1.1* | 16 | 13 | 0.890 | 1.33 (1 – 4) |

For the *cpmp, cpp* and *csp* microhaplotypes, a direct comparison between nanopore and Illumina sequencing was possible because the shorter amplicons used for nanopore sequencing were within the longer amplicons previously sequenced by Illumina ^13^. We extracted the respective regions of the microhaplotype markers used for nanopore sequencing to directly compare them. Overall, 96.8% (30/31) haplotypes were concordant with both technologies for *cpmp* (Figure 4a), 94.7% (18/19) for *cpp* (Figure 4b), and 86.3% (19/22) for *csp* (Figure 4c). We further investigated the additional discriminatory power when the entire, longer amplicons (∼300-450 bp ^11,12^) using Illumina sequencing were used compared to the nanopore AmpSeq panel of shorter amplicons (∼150-200 bp). Longer amplicons should theoretically be more diverse as they capture additional SNPs, increasing the number of haplotypes. For *cpmp*, 93.8% (30/32) haplotypes were found using both technologies (Supplementary Figure 5a). For *cpp*, the longer amplicon identified six more haplotypes with Illumina, and only 75% (18/24) were distinguished by the shorter amplicon using nanopore AmpSeq (Supplementary Figure 5b). For *csp*, there was high concordance, with 95% (19/20) of haplotypes also found using nanopore AmpSeq (Supplementary Figure 5c), and the shorter amplicon even detected two additional haplotypes missed by Illumina, possibly due to higher PCR efficiency of shorter amplicons. Overall, the high concordance between the two methods supports multiplexed nanopore AmpSeq as a reliable method for identifying and distinguishing different parasite clones.

**Figure 4.**
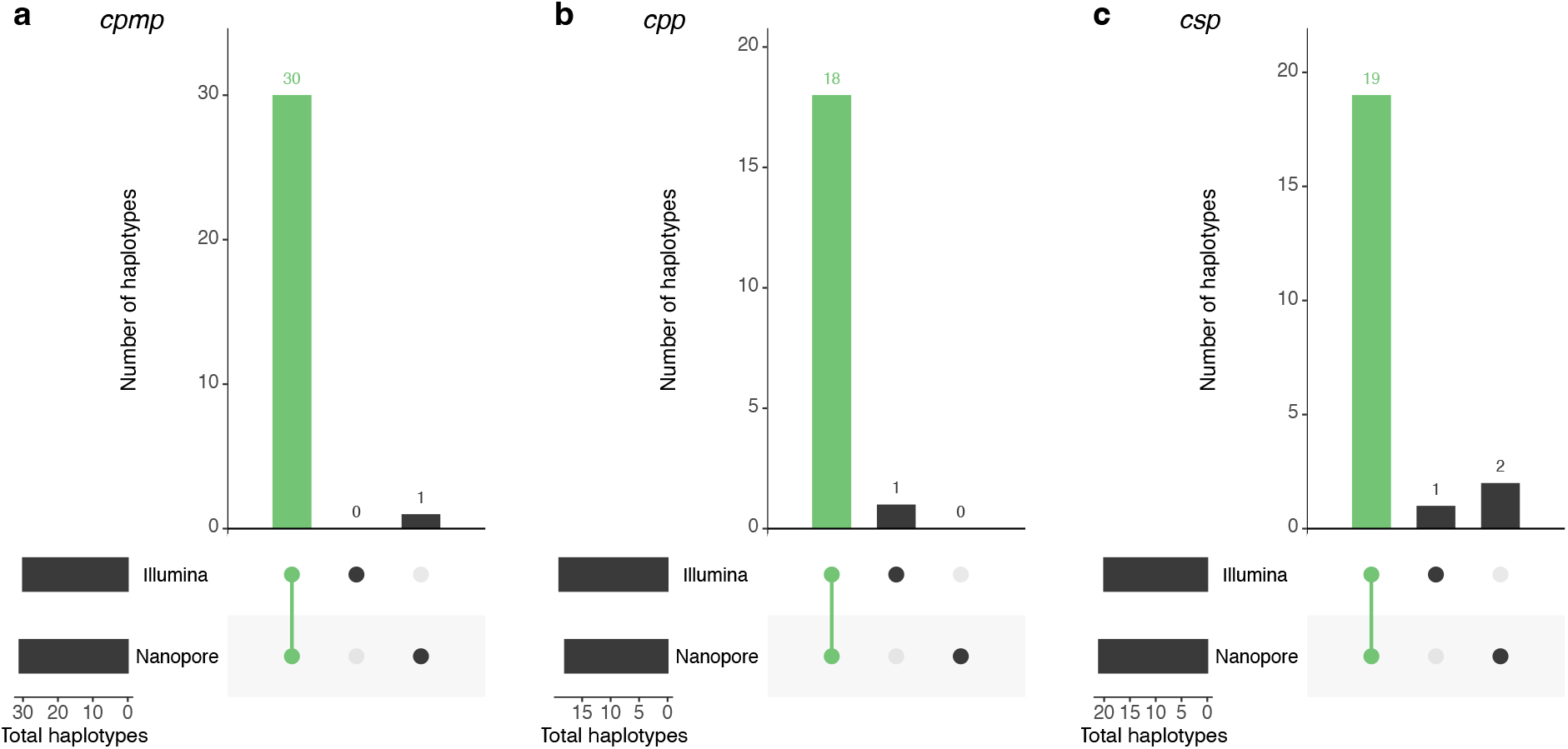
Number of haplotypes of three different marker between nanopore and Illumina sequencing. UpSetR plots showing the numbers of haplotypes shared between the two different sequencing technologies and different amplicons for the genes (**a**) *cpmp* (PF3D7_0104100), (**b**) *cpp* (PF3D7_1475800), and (**c**) *csp* (PF3D7_0304600). Haplotypes that were found with both methods are highlighted in green. Sequences from Illumina sequencing were trimmed to the same length of markers used for nanopore sequencing.

### Distinguishing recrudescence from new infection in an antimalarial drug clinical trial

The ability and reliability of nanopore AmpSeq to distinguish recrudescence from new infection were assessed using 20 paired patient samples. Concordance among the six microhaplotype markers was high, with all markers providing the same outcome for 90% (18/20) of the patients (Table 2). Recrudescence and new infection outcomes were also compared to AmpSeq data obtained from Illumina sequencing (from ref.^13^), with almost identical results between the two sequencing technologies (Table 2). The final genotyping outcome was determined using two different algorithms: the WHO/MMV algorithm and the 2/3 algorithm (Table 3). The outcomes for nanopore AmpSeq using both algorithms were concordant for all 20 patients, and agreement with genotyping outcomes from Illumina AmpSeq was 100%.

**Table 2.**
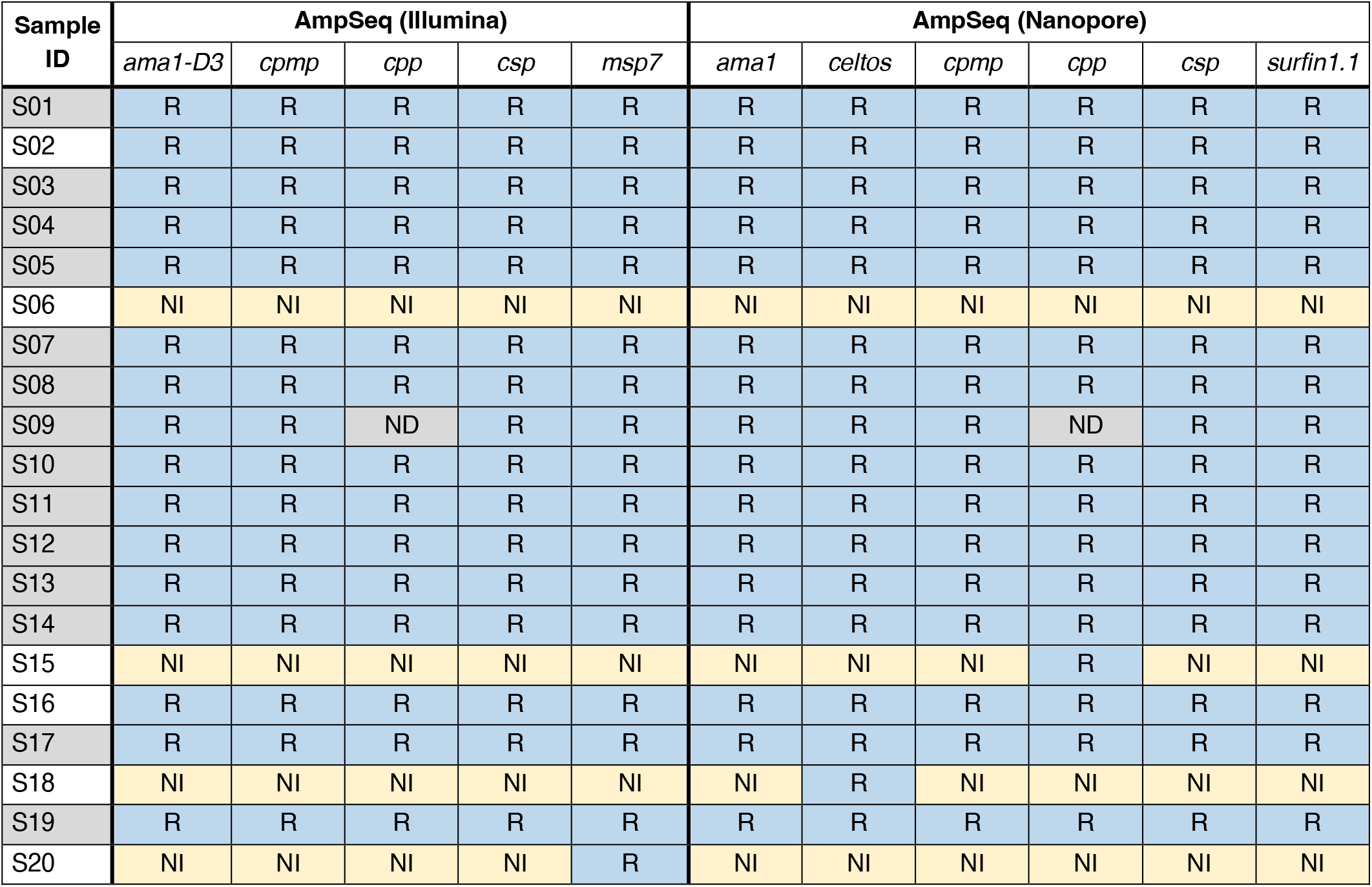
New infection and recrudescence outcome of AmpSeq using Illumina and nanopore sequencing using 20 paired patient samples. R = recrudescence outcome (blue shading); NI = new infection outcome (yellow shading); ND = not determined. Samples shaded in grey represent recurrent samples with the *PfATP4* G538S mutation associated with resistance to cipargamin. Illumina data were previously generated in ref ^*13*^.

**Table 3.** Results of PCR-correction outcomes determined by AmpSeq using Illumina and nanopore of 20 paired pre- and post-treatment samples. Both, the WHO/MMV and 2/3 algorithm (4/5 for Illumina and 4/6 for nanopore) were used. Samples shaded in grey represent recurrent samples with the *PfATP4* G538S mutation associated with resistance to cipargamin. Illumina data were previously generated in ref ^*13*^.

| Sample ID | AmpSeq Illumina (*ama1-D3, cpmp, cpp, csp, msp7*) |  | AmpSeq Nanopore (*ama1, celtos, cpmp, cpp, csp, surfin1.1*) |  |
| --- | --- | --- | --- | --- |
|  | WHO/MMV | 2/3 | WHO/MMV | 2/3 |
| S01 | R | R | R | R |
| S02 | R | R | R | R |
| S03 | R | R | R | R |
| S04 | R | R | R | R |
| S05 | R | R | R | R |
| S06 | NI | NI | NI | NI |
| S07 | R | R | R | R |
| S08 | R | R | R | R |
| S09 | R | R | R | R |
| S10 | R | R | R | R |
| S11 | R | R | R | R |
| S12 | R | R | R | R |
| S13 | R | R | R | R |
| S14 | R | R | R | R |
| S15 | NI | NI | NI | NI |
| S16 | R | R | R | R |
| S17 | R | R | R | R |
| S18 | NI | NI | NI | NI |
| S19 | R | R | R | R |
| S20 | NI | NI | NI | NI |
| **Total number of recrudescence's** | **16** | **16** | **16** | **16** |

Finally, we estimated pairwise genetic relatedness for each of the 20 paired patient samples (baseline and recurrent sample) using identity-by-descent (IBD) ^20^. As expected, the four samples classified as new infections showed no genetic relatedness (IBD<0.1), whereas the 16 sample pairs classified as recrudescence showed high genetic relatedness (IBD=1, Supplementary Table 7). Using pairwise IBD confirmed the outcomes of recrudescence and new infection, as high relatedness is expected in sample pairs with recrudescent infections and low to no relatedness in pairs with new infections.

## DISCUSSION

PCR-correction is crucial for accurate estimation of antimalarial drug efficacy. Genotyping assays must be highly sensitive to detect minority clones in polyclonal infections, reproducible, and utilize markers with high diversity and low allelic frequencies ^9^. Previous studies have shown that AmpSeq provides good sensitivity and reproducibility, but also higher concordance between markers compared to other genotyping methods. Furthermore, using three to five highly diverse amplicons was sufficient to reliably distinguish between recrudescence and new infections, supporting the adoption of AmpSeq as the standard for PCR-correction ^13,19^. However, many malaria-endemic countries currently lack the necessary NGS capacity for AmpSeq. Smaller labs often ship samples internationally for sequencing, which delays results. While capacity is gradually improving, there is a need for efficient, cost-effective, reliable, and accessible tools to strengthen genomic capabilities in research and public health institutions in malaria-endemic countries before AmpSeq can be widely adopted for routine TES.

In this study, we optimized a multiplexed nanopore AmpSeq of *P. falciparum* microhaplotypes. Our nanopore AmpSeq panel can accurately estimate within-host diversity and is potentially well-suited for distinguishing recrudescence from new infection in clinical trials. The protocol, from DNA extraction to results, takes about three to four days. Using laboratory strain mixtures and paired clinical samples, our assay demonstrated high and uniform coverage across microhaplotypes, crucial for accurate variant calling, particularly in identifying minority clones. The method is highly sensitive and reproducible, providing data for low parasite density samples and detecting minor alleles in polyclonal infections at a WSAF as low as 1.9% with good specificity. The detection of minority clones at concentrations up to 50-fold lower is comparable to other *P. falciparum* AmpSeq panels designed for Illumina platforms ^14–16^. After removing reads below 99% accuracy (≥Q20) and applying a WSAF threshold of ≥1%, results were highly specific, with no false-positive haplotypes detected. To fully exploit the information content from diverse microhaplotype loci, especially in polyclonal infections, it is essential to use haplotype inference tools capable of accurately identifying complex *P. falciparum* infections with unknown haplotype proportions. While such methods exist for short-read Illumina data, only our previously developed workflow could capture all information from polyclonal infections using nanopore data ^22^. For our workflow, we adapted existing short-read haplotype inference tools *HaplotypR* ^11^ and *DADA2* ^26^, using only high-quality reads (≥Q20) and appropriate cut-offs. Given the short length of the amplicons, only few reads have sequencing errors, allowing for effective haplotype inference since distinguishing true variation from errors is possible, as supported by our results. However, this approach is currently not suitable for longer amplicons due to the high accumulation of errors, even after filtering to ≥Q20, making it challenging to differentiate between true variants and sequencing errors.

Our panel showed high genetic diversity and low allelic frequencies in the patients’ samples, with up to 31 different haplotypes for *cpmp*, providing high discriminatory power. However, two of the six markers used in our panel were not able to distinguish two well-characterized laboratory strains (FCB1 and K1 for *ama1*; FCB1 and HB3 for *csp*). These results are intriguing as both markers have good diversity, with H_E_ ≥ 0.95, and a marker with lower diversity, i.e. *surfin1*.*1* (H_E_= 0.89) was able to distinguish all the four laboratory strains used. Moreover, *ama1* and *csp* had concordant results with other markers in distinguishing recrudescence from new infections in patients’ samples. These findings highlight the need for thorough investigation to validate new techniques and new markers using both laboratory strains to assess sensitivity and specificity of the assay and patients’ samples for diversity and allelic frequency of the markers used. Indeed the optimal combination of assay and markers will need to have a good balance between sensitivity, specificity of the assay and genetic diversity of the markers used.

Previous studies have shown that PCR-correction outcomes can vary widely depending on the algorithm used ^13,19^. The six markers used in this study consistently gave concordant results in 90% of samples analyzed, and more importantly, the final PCR-corrected outcomes were 100% identical for all samples, independent of the algorithm used to analyze the data. Further, the outcomes were also identical to those previously obtained with longer amplicons sequenced on the Illumina platform for the same samples ^13^. This underscores that nanopore AmpSeq is a reliable and comparable alternative to Illumina AmpSeq. Additionally, a relatedness-based approach (i.e., IBD) provided the same results as the 2/3 and WHO/MMV algorithms, explicitly accounting for population allele and providing uncertainty measures ^20^. Such novel approaches could potentially provide even more robust estimates of interhost relatedness by using a larger set of microhaplotypes and accounting for missing alleles, thus providing a more accurate final result.

ONT devices are a relatively cheaper alternative to traditional sequencing methods, offering rapid data generation and enabling in-country sequencing, thus minimizing result turnaround times. Despite its apparent benefits and continuous improvements in sequencing performance ^29^, nanopore sequencing of *P. falciparum* remains underutilized ^22–24,30^. While previous studies have demonstrated the feasibility of using nanopore AmpSeq for various use cases ^22–24,30^, including sequencing in malaria-endemic countries ^22,23^, this is the first study to use nanopore sequencing to distinguish recrudescence from new infection in antimalarial drug efficacy studies. Despite the ability to perform long-read AmpSeq with nanopore-based sequencing, we chose to use short amplicons to distinguish recrudescence from new infection for several reasons. First, shorter amplicons typically have higher amplification efficiency compared to longer alternatives. Second, targeting short amplicons allows to focus on the most informative regions and avoid error-prone homopolymers or tandem repeats. Indeed, nanopore sequencing struggles to accurately resolve low-complexity regions ^24,31^ that are abundant in the *P. falciparum* genome ^32^. In addition, previous studies using longer amplicons of the highly polymorphic *ama1, csp, msp1*, and *msp2* genes for nanopore AmpSeq were unable to infer individual haplotypes or detect minority clones in polyclonal infections ^23,24^.

Our study has several limitations. First, for two markers, *ama1* and *csp*, the LOD could only be determined for two out of four strains because of identical alleles. Second, we did not assess inter-assay reproducibility, either within the same laboratory or between laboratories. Third, the small number of patient samples, with a high proportion of recrudescence, limits the generalizability of the results. Lastly, our assay was designed to be an easy-to-use approach achieving high and uniform coverage. Thus, we chose to not transfer the previously used Illumina AmpSeq protocol ^13^ to the ONT platform, but rather to use a multiplexed PCR of short microhaplotypes with high amplification efficiency. Although some genes are identical, a direct comparison with the protocol using Illumina was only possible to a limited extent.

In conclusion, our study demonstrates the suitability of nanopore AmpSeq for genotyping *P. falciparum* to distinguish recrudescence from new infection. We show that detecting minority clones in polyclonal infections and characterizing within-host diversity is feasible using nanopore AmpSeq targeting short, highly diverse microhaplotypes. However, further studies with different laboratory strains and larger clinical sample sizes are needed to confirm the specificity of the assay in detecting different parasite clones and its capability to correctly distinguish recrudescence from new infection compared to Illumina AmpSeq. ONT sequencing platforms are affordable, portable, scalable, and well-suited for resource-limited endemic settings, offering a promising solution for in-country sequencing and rapid PCR-corrected estimates of drug failure using AmpSeq.

## DATA AVAILABILITY

Raw sequence data is available for download at the NCBI Sequence Read Archive (Accession numbers: pending). All sequence data and code used in this analysis can be found at (link Zenodo repository. All analyses were completed in R version 4.1.2.

## CODE AVAILABILITY

All analyses were completed in R version 4.1.2 using published R packages, as described in the methods section. Code used in this analysis can be found at (link Zenodo repository). The source code for software *HaplotypR*, including a tutorial with example datasets, is available at https://github.com/lerch-a/HaplotypR.

## ACKNOWLEDGEMENTS

We would like to thank Annina Schnoz from Swiss Tropical and Public Health Institute for providing the laboratory strain mixtures.

## AUTHOR INFORMATION

### Contributions

A.H.: Conceptualization, Methodology, Investigation, Formal analysis, Visualization, Writing - Original Draft. A.L.: Investigation, Software, Writing - Review & Editing. C.N.: Supervision, Funding acquisition, Resources, Writing - Review & Editing. All authors reviewed and approved the final version of this manuscript.

