## Supplementary_Information for "Multiplexed nanopore amplicon sequencing to distinguish recrudescence from new infection in antimalarial drug trials"

Supplementary information for the paper Holzschuh et al. ***Multiplexed nanopore amplicon sequencing to distinguish recrudescence from new infection in antimalarial drug trials***

TABLE OF CONTENTS


1. Supplementary Table 1 Page 2
2. Supplementary Table 2 Page 3
3. Supplementary Table 3 Page 4
4. Supplementary Table 4 Page 5
5. Supplementary Table 5 Page 6
6. Supplementary Table 6 Page 7
7. Supplementary Table 7 Page 8

8. Supplementary Figure 1 Page 9
9. Supplementary Figure 2 Page 10
10. Supplementary Figure 3 Page 11
11. Supplementary Figure 4 Page 12
12. Supplementary Figure 5 Page 13

**Supplementary Table 1. Laboratory strain mixtures at different ratios.** Mixtures were prepared and used previously [1].

| Sample ID | Strain ratio | | | | Parasites/$\mu$L blood | | | | WSAF | | | |
|---|---|---|---|---|---|---|---|---|---|---|---|---|
| | 3D7 | K1 | HB3 | FCB 1 | 3D7 | K1 | HB3 | FCB 1 | 3D7 | K1 | HB3 | FCB 1 |
| S01_new | 1 | 0 | 0 | 0 | 1,000 | 0 | 0 | 0 | 1 | 0 | 0 | 0 |
| S02_new | 0 | 1 | 0 | 0 | 0 | 1,000 | 0 | 0 | 0 | 1 | 0 | 0 |
| S03_new | 0 | 0 | 1 | 0 | 0 | 0 | 1,000 | 0 | 0 | 0 | 1 | 0 |
| S04_new | 0 | 0 | 0 | 1 | 0 | 0 | 0 | 1,000 | 0 | 0 | 0 | 1 |
| S05_new | 1 | 1 | 1 | 1 | 1,000 | 1,000 | 1,000 | 1,000 | 0.25 | 0.25 | 0.25 | 0.25 |
| S05 | 1 | 1 | 1 | 1 | 10 | 10 | 10 | 10 | 0.25 | 0.25 | 0.25 | 0.25 |
| S06 | 5 | 1 | 1 | 1 | 50 | 10 | 10 | 10 | 0.625 | 0.125 | 0.125 | 0.125 |
| S07 | 1 | 5 | 1 | 5 | 10 | 50 | 10 | 50 | 0.083 | 0.417 | 0.083 | 0.417 |
| S08 | 1 | 5 | 5 | 5 | 10 | 50 | 50 | 50 | 0.063 | 0.313 | 0.313 | 0.313 |
| S09 | 1 | 10 | 1 | 1 | 10 | 10 | 10 | 10 | 0.077 | 0.769 | 0.077 | 0.077 |
| S10 | 1 | 1 | 10 | 10 | 10 | 10 | 100 | 100 | 0.045 | 0.045 | 0.455 | 0.455 |
| S11 | 10 | 1 | 10 | 10 | 100 | 10 | 100 | 100 | 0.323 | 0.032 | 0.323 | 0.323 |
| S12 | 1 | 1 | 15 | 1 | 10 | 10 | 150 | 10 | 0.056 | 0.056 | 0.833 | 0.056 |
| S13 | 15 | 15 | 1 | 1 | 150 | 150 | 10 | 10 | 0.469 | 0.469 | 0.031 | 0.031 |
| S14 | 15 | 15 | 1 | 15 | 150 | 150 | 10 | 150 | 0.326 | 0.326 | 0.022 | 0.326 |
| S15 | 1 | 1 | 1 | 20 | 10 | 10 | 10 | 200 | 0.043 | 0.043 | 0.043 | 0.87 |
| S16 | 20 | 1 | 20 | 1 | 200 | 10 | 200 | 10 | 0.476 | 0.024 | 0.476 | 0.024 |
| S17 | 20 | 20 | 20 | 1 | 200 | 200 | 200 | 10 | 0.328 | 0.328 | 0.328 | 0.016 |
| S18 | 50 | 1 | 1 | 1 | 500 | 10 | 10 | 10 | 0.943 | 0.019 | 0.019 | 0.019 |
| S19 | 1 | 50 | 1 | 50 | 10 | 500 | 10 | 500 | 0.01 | 0.490 | 0.01 | 0.490 |
| S20 | 1 | 50 | 50 | 50 | 10 | 500 | 500 | 500 | 0.007 | 0.331 | 0.331 | 0.331 |
| S21 | 100 | 1 | 1 | 1 | 1000 | 10 | 10 | 10 | 0.971 | 0.01 | 0.01 | 0.01 |
| S22 | 1 | 100 | 1 | 100 | 10 | 1000 | 10 | 1000 | 0.005 | 0.495 | 0.005 | 0.495 |
| S23 | 1 | 100 | 100 | 100 | 10 | 1000 | 1000 | 1000 | 0.003 | 0.332 | 0.332 | 0.332 |

**Table 2. *P. falciparum* highly polymorphic microhaplotypes targeted by the 6-plex amplicon panel.** Median length across all amplicons is 231 bp (range: 179-250).

| Gene name (*PlasmoDB* Gene ID) | Chromosome | Start | End | Amplicon size (including primer) | Associated phenotype |
|---|---|---|---|---|---|
| Apical membrane antigen 1, *ama1* (PF3D7_1133400) | Pf3D7_11_v3 | 1294271 | 1294520 | 250 bp | Vaccine candidate antigen; potential for use as a marker of complexity of infection |
| Cell traversal protein for ookinetes and sporozoites, *celtos* (PF3D7_1216600) | Pf3D7_12_v3 | 659859 | 660037 | 179 bp | Vaccine candidate antigen, potential for use as a marker of complexity of infection |
| Conserved Plasmodium membrane protein, *cpmp* (PF3D7_0104100) | Pf3D7_01_v3 | 180130 | 180370 | 241 bp | Potential for use as a marker of complexity of infection |
| Conserved Plasmodium protein, *cpp* (PF3D7_1475800) | Pf3D7_14_v3 | 3121036 | 3121267 | 232 bp | Potential for use as a marker of complexity of infection |
| Circumsporozoite protein, *csp* (PF3D7_0304600) | Pf3D7_03_v3 | 221468 | 221655 | 188 bp | Leading vaccine and monoclonal antibody target antigen; potential for use as a marker of complexity of infection |
| Surface-associated interspersed protein 1.1, *surfin1.1* (PF3D7_0113100) | Pf3D7_01_v3 | 495942 | 496171 | 230 bp | Potential for use as a marker of complexity of infection |

**Supplementary Table 3. Primer sequences for the 6-plex microhaplotype panel.**

| Primer | Direction | Sequence | Source | Stock concentration (mM) | Concentration in final pool (μM) | Final concentration in PCR reaction (μM) |
|---|---|---|---|---|---|---|
| ***ama1*** | Forward | GAACTCAATATAGACTTCCATCAGG | [2] | 1 | 7 | 0.14 |
| | Reverse | CCTGCATGTCTTGAACATAAAGTC | [2] | 1 | 7 | 0.14 |
| ***celtos*** | Forward | CTGGTACTATTATACCATATGTTGC | [2] (adapted from [3]) | 1 | 15 | 0.3 |
| | Reverse | TCACCAACCTTTTTAGAATCAAGC | [2] (adapted from [3]) | 1 | 15 | 0.3 |
| ***cpmp*** | Forward | GGAAGCTATAGGTATCAGATCC | [2] | 1 | 10 | 0.2 |
| | Reverse | TAGAATACGTGCTTTATAAACAAAGAG | [2] | 1 | 10 | 0.2 |
| ***cpp*** | Forward | AACACAATCTTCCTTAGCCAATTC | [2] | 1 | 8 | 0.16 |
| | Reverse | ATTACTACCTTTCAGCATATCCGA | [2] | 1 | 8 | 0.16 |
| ***csp*** | Forward | GACCCAAACCGAAATGTAGATG | [2] | 1 | 10 | 0.2 |
| | Reverse | GAGCCAGGCTTTATTCTAACTTG | [2] | 1 | 10 | 0.2 |
| ***surfin1.1*** | Forward | CACCAAAATATTATATACCACAAGAC | [2] (adapted from [3]) | 1 | 15 | 0.3 |
| | Reverse | GGAAAATCTTTGGTGGGAAAAATAG | [2] (adapted from [3]) | 1 | 15 | 0.3 |

**Supplementary Table 4. Thermal cycling conditions of 6-plex microhaplotype panel.**

| Cycling step | Temp (°C) | Time | # of cycles |
|---|---|---|---|
| Initial denaturation | 95 | 3 min | 1 |
| Denaturation | 98 | 15 sec | 35 |
| Annealing | 56 | 15 sec | |
| Extension | 72 | 30 sec | |
| Final Extension | 72 | 2 min | 1 |
| Hold | 4 | ∞ | |

Note: Use a heated lid set to 105°C and set the sample volume to 25 µL.

**Supplementary Table 5. ONT sequencing run characteristics.**

| Run | ONT chemistry used | Samples per run (of which controls) | Run time | Total reads | Mean Q-score (accuracy)* | Median Q-score (accuracy)* | Reads with ≥Q20, pass (%) |
|---|---|---|---|---|---|---|---|
| Run1 (Control mixtures) | Kit 14, R10.4.1 | 75 (3) | 23h 30min | 11,151,393 | 16.2 (97.6%) | 21.6 (99.3%) | 6,927,660 (62.1%) |
| Run2 (Paired patient samples) | Kit 14, R10.4.1 | 44 (4) | 18h 10min | 6,510,788 | 15.6 (97.2%) | 20.9 (99.2%) | 3,691,523 (56.7%) |

* With *dorado v0.7.0* using the super-accurate (sup) model (dna_r10.4.1_e8.2_400bps_sup@v5.0.0).

**Supplementary Table 6. Different primer concentrations tested for primer balancing.** Primer pools ‘old’, ‘new1, ‘new2, and ‘new3’ were tested at annealing temperatures 56 °C and 58 °C. 56 °C was found to be superior and ultimately in a second sequencing round, primer pool ‘new4’ (highlighted) was found to produce most even coverage across all amplicons.

| **Amplicon** | **Primer concentrations in different pools ($\mu$M)** | | | | | |
|---|---|---|---|---|---|---|
| | old* | new1 | new2 | new3 | **new4** | new5 |
| *ama1* | 5 | 15 | 15 | 10 | **7** | 8 |
| *celtos* | 10 | 10 | 10 | 10 | **15** | 12 |
| *cpmp* | 10 | 15 | 10 | 10 | **10** | 10 |
| *cpp* | 5 | 20 | 15 | 10 | **8** | 9 |
| *csp* | 5 | 5 | 5 | 10 | **10** | 10 |
| *surfin1.1* | 15 | 10 | 10 | 10 | **15** | 12 |

*Primer pool from [2].

**Supplementary Table 7. Pairwise IBD for the 20 paired patient samples using nanopore AmpSeq.** IBD was estimated using haplotypes from the six microhaplotype nanopore AmpSeq data. Pairwise *P* values were determined by likelihood-ratio adjusted for 1-sided tests implemented in *dcifer* [3].

| Sample ID | Pairwise IBD | *P* value |
|---|---|---|
| **S01** | 1 | 2.74e-09 |
| **S02** | 1 | 2.15e-06 |
| **S03** | 1 | 4.22e-09 |
| **S04** | 1 | 0.00013 |
| **S05** | 1 | 8.11e-10 |
| **S06** | 0 | >0.05 |
| **S07** | 1 | 5.64e-09 |
| **S08** | 1 | 3.19e-08 |
| **S09** | 1 | 9.72e-08 |
| **S10** | 1 | 5.94e-09 |
| **S11** | 1 | 4.73e-10 |
| **S12** | 1 | 2.95e-06 |
| **S13** | 1 | 1.49e-07 |
| **S14** | 1 | 1.09e-06 |
| **S15** | 0.09 | 0.25635 |
| **S16** | 1 | 1.14e-07 |
| **S17** | 1 | 1.07e-09 |
| **S18** | 0 | >0.05 |
| **S19** | 1 | 3.17e-07 |
| **S20** | 0 | >0.05 |

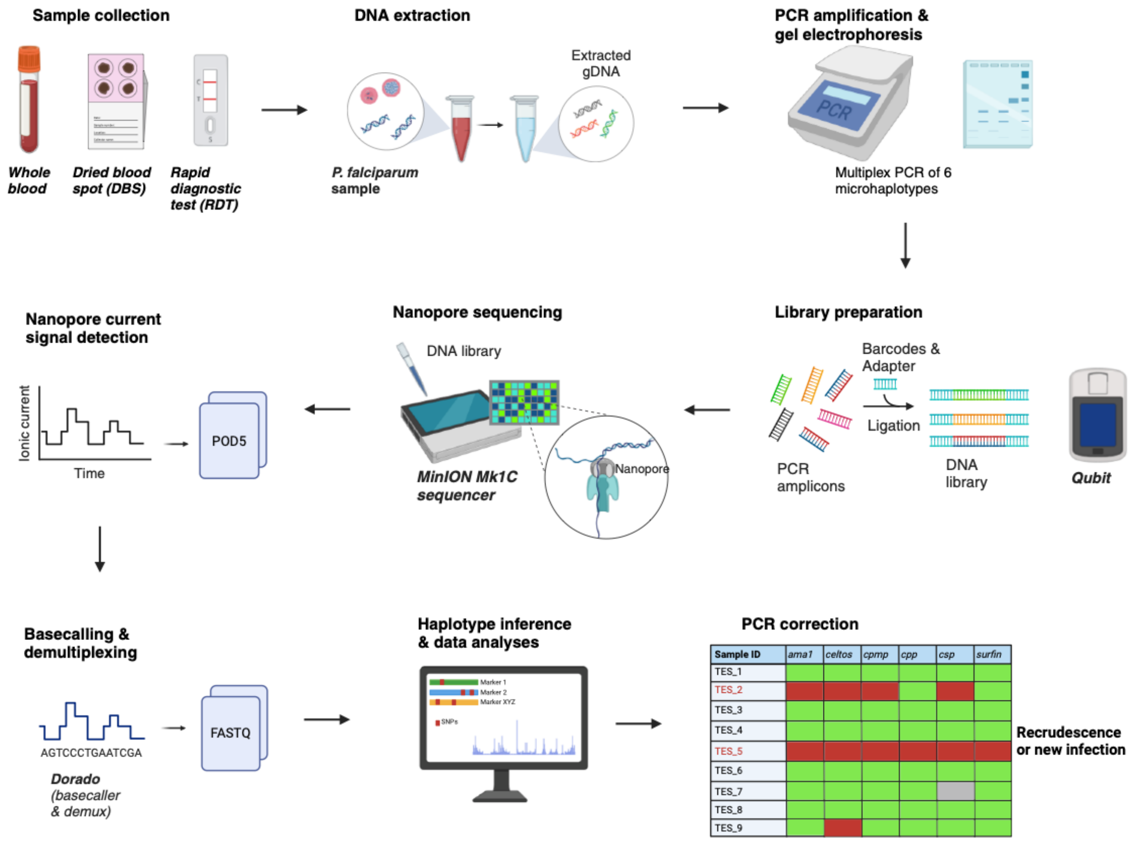


**Supplementary Figure 1. Overview of the *P. falciparum* multiplexed nanopore AmpSeq approach using Oxford Nanopore Technologies (ONT).** DBS and RDT samples have been successfully sequenced using the 6-plex microhaplotype panel, however this is dependent on parasite density [2]. The MinION Mk1B device could also be used instead of the MinION Mk1C. Created with BioRender.com.

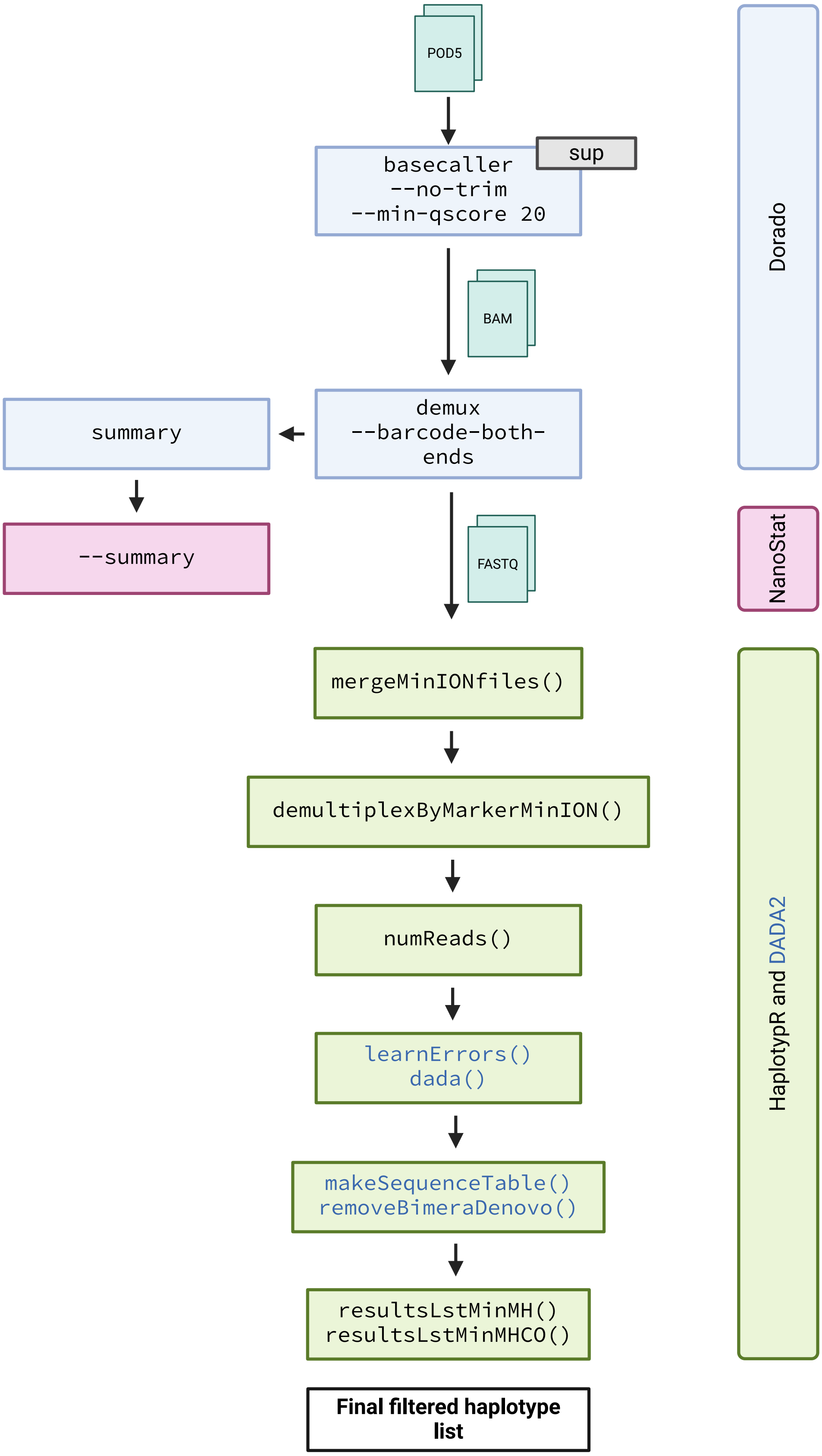


**Supplementary Figure 2. Overview of the bioinformatics workflow for haplotype inference of microhaplotypes using ONT sequencing data.** Different software and packages used are shown on the right. Created with BioRender.com.

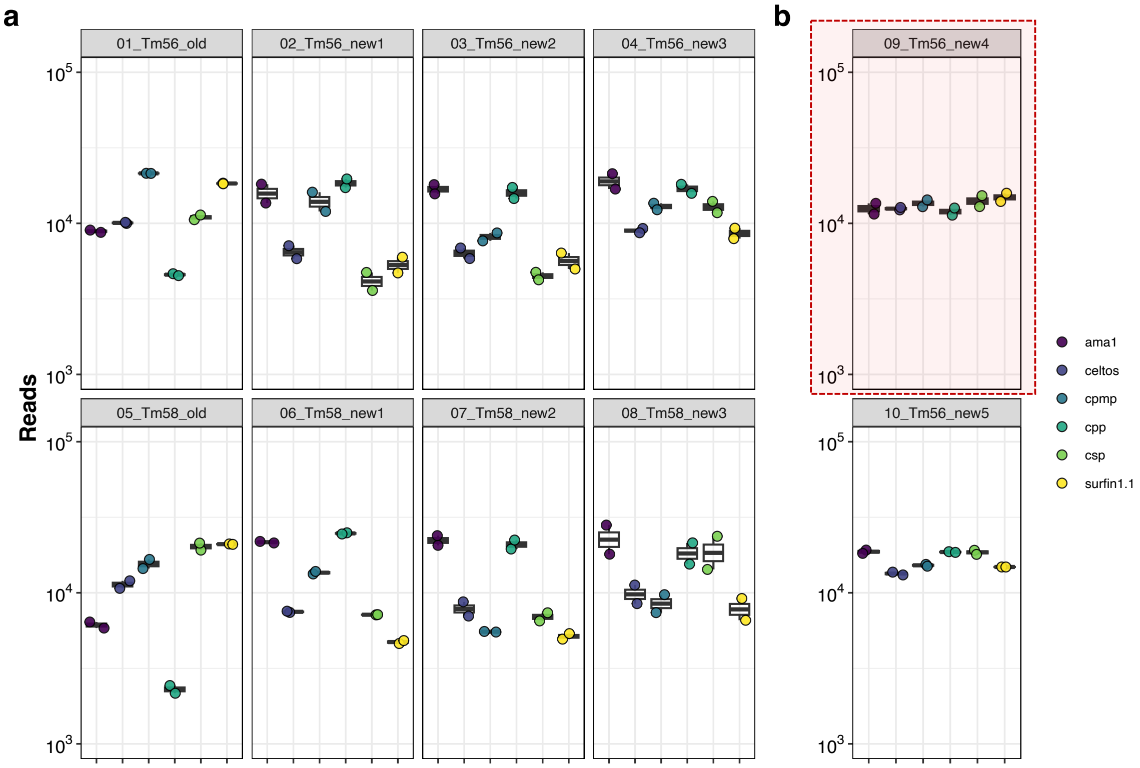


**Supplementary Figure 3. Coverage profile of target amplicons using different primer concentrations and two PCR annealing temperatures.** (**a**) First primer-balancing sequencing run using 8 different conditions (4 different primer pool concentrations and 2 different annealing temperatures). (**b**) Second primer-balancing sequencing run using two additional primer pool concentrations, both at 56 ºC annealing temperature. Condition 09 was selected (primer pool new4), highlighted in red, showing most even coverage across all 6 amplicons. All conditions were tested in duplicate using *P. falciparum* strain 3D7 at 1,000p/µL. Primer concentrations of different pools are shown in Supplementary Table 6.

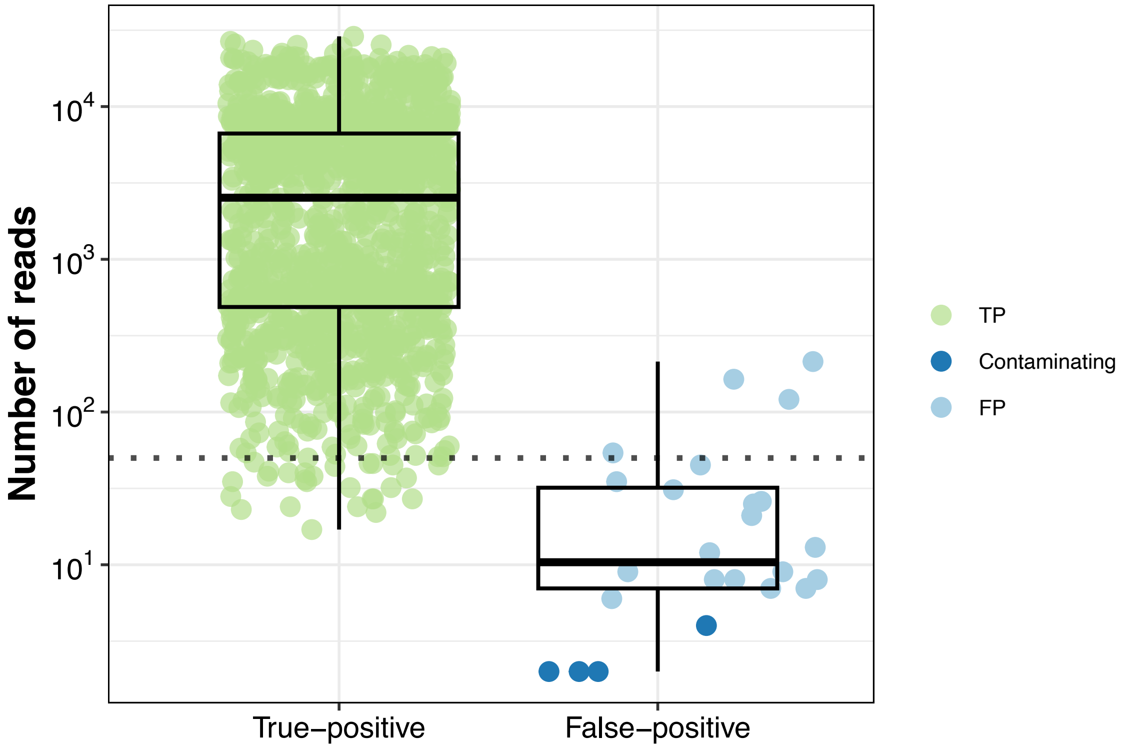


**Supplementary Figure 4. Number of reads for true-positive and false-positive haplotypes.** Only 4 incorrect haplotypes were above the cut-off of ≥50 reads per haplotype (dotted line). The majority of incorrect reads were false-positive haplotypes (i.e., amplification or sequencing errors) and not contaminating haplotypes (i.e., true-positive haplotypes originating from any other *P. falciparum* strain present in another sample). Importantly, all false-positive haplotypes fall below the within sample frequency cut-off of ≥1%. 26/1392 (1.9%) TP haplotypes fall below the ≥50 reads per haplotype cut-off. The dotted line indicates the minimum coverage of ≥50 reads per haplotype. Note: no cut-off was applied to the data for haplotype inference, meaning all haplotypes (i.e., all reads) identified are shown.

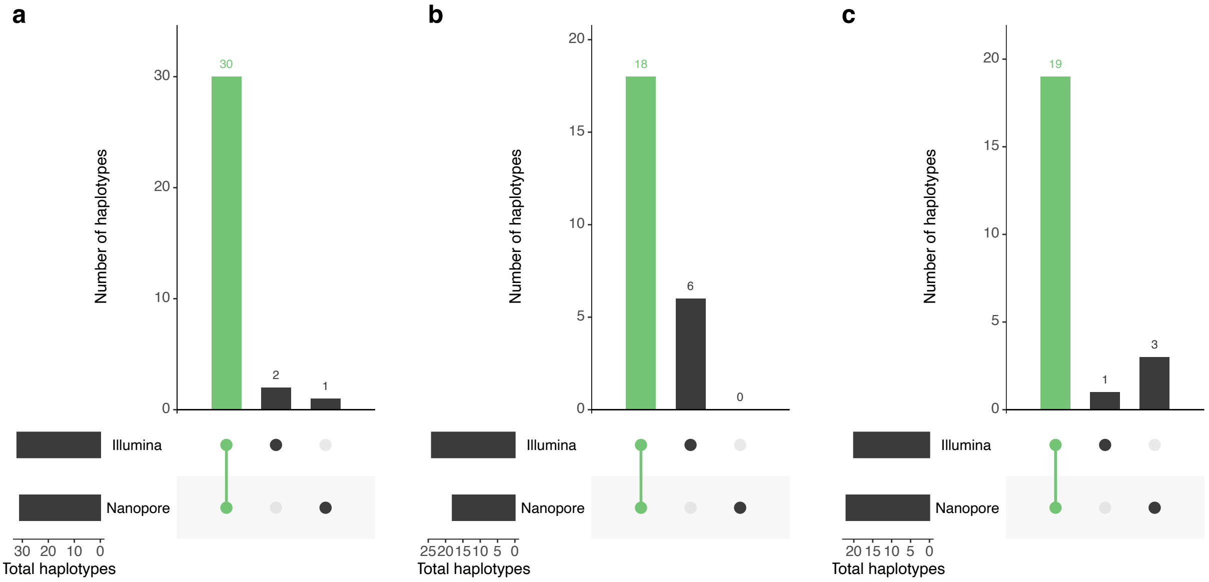


**Supplementary Figure 5. Number of haplotypes of three different marker between nanopore and Illumina sequencing.** UpSetR plots showing the numbers of haplotypes shared between the two different sequencing technologies and different amplicons for the genes (**a**) *cpmp* (PF3D7_0104100), (**b**) *cpp* (PF3D7_1475800), and (**c**) *csp* (PF3D7_0304600). Haplotypes that were found with both methods are highlighted in green.
